# Genome analysis of *Legionella pneumophila* ST901 an Italian endemic strain

**DOI:** 10.1101/2023.06.08.544189

**Authors:** Maria Luisa Ricci, Silvia Fillo, Francesco Giordani, Andrea Ciammaruconi, Antonietta Girolamo, Anna Anselmo, Anella Monte, Valerio Cusimano, Maria Grazia Caporali, Maria Cristina Rota, Markus Petzol, Baharak Afshar, Florigio Lista, Christian Luck, Maria Scaturro

**Affiliations:** Department of Infectious Diseases, Istituto Superiore di Sanità, Rome Italy; ESCMID Study Group for Legionella Infections (ESGLI), Basel, Switzerland; Defense Institute for Biomedical Sciences, Rome, Italy; Italian National Research Council - Institute for Systems Analysis and Computer Science, Rome, Italy; Institut für Medizinische Mikrobiologie und Virologie, Fiedlerstr. 42 -01307 Dresden; Respiratory and Vaccine Preventable Bacteria Reference Unit (RVPBRU), International Health Regulations Strengthening Project Clinical Scientist, UK Health Security Agency

**Keywords:** cgMLST, CRISPR-cas system, Legionella pneumophila, Pangenome, SNPs, ST901

## Abstract

*Legionella pneumophila* typing, based on SBT analysis, has been allowing to identify sequence types (ST) responsible for many Legionnaires’ disease cases. ST901 has so far been identified only in a town of the northern Italy, where it caused numerous travel-associated Legionnaires’ disease cases over a 32-year period. In this study, ST901 were analyzed by whole genome sequencing in looking for both differences and similarities inside this ST and with other unrelated *Legionella* strains. To this aim core genome multi locus sequence type, single nucleotide polymorphisms and pangenome analyses were performed. ST901 resulted highly similar each other, characterized by I-C CRISPR-cas system, located on the chromosome and by a putative plasmid, highly similar to that found in Lp strain Lens, which contains I-F CRISPR-cas system. Accessory genomic islands, either already described or specifically found in ST901, were also shown.

In conclusion ST901 has been quite preserved over the time. The co-occurrence of two phage resistance systems and genomic islands of virulence could have made the ST901 enough virulent to cause such a high number of cases.

**Summary blurb:** ST901 appears as endemic of the Northern Italy, with virulence traits similar to those in Alcoy and Corby, and with two CRISPR-cas systems which probably preserved them in their environmental niche.

## Introduction

*Legionella* is a widespread genus in fresh water habitats where it lives in close association with free-living amoebae feeding on biofilm. To date 63 *Legionella* species and 3 subspecies (https://lpsn.dsmz.de/genus/legionella) [1] are known and *Legionella pneumophila* is the most commonly detected species in human illness cases. Although 15 serogroups have been identified serogroup (sg) 1 is responsible for 83% of infections in Europe [2]. In 2020 in Europe the notification rate varied from 0.5 to 5.7 cases per 100,000 population, and four countries, France, Germany, Italy and Spain, accounted for 72% of all notified cases [2]. In US the reported incidence also increased more than three-fold in the last 10 years [3]. Most cases are both sporadic and community-acquired but a fair number of cases are travel-associated Legionnaires’ disease (TALD). However, the disease remains underestimated and in Italy a different incidence between Southern and Northern regions is reported, with 10.5 cases per million population in the Southern compared to 50 cases per million population in the Northern regions [4]. In respect of the TALD, considerable attention is paid to Italy that still remains the country with the highest number of TALD clusters reported at the European Legionnaires’ Disease Surveillance Network, often involving reoffending accommodation sites, in spite of the implemented control and prevention measures [2-4].

Since 2004, according to the Italian National surveillance register, 61 TALD cases occurred in different hotels in Molveno, a small (35 Km^2^) town with a high tourist vocation of the Northern Italy, with many hotels overlooking the homonymous lake (Figure 1).

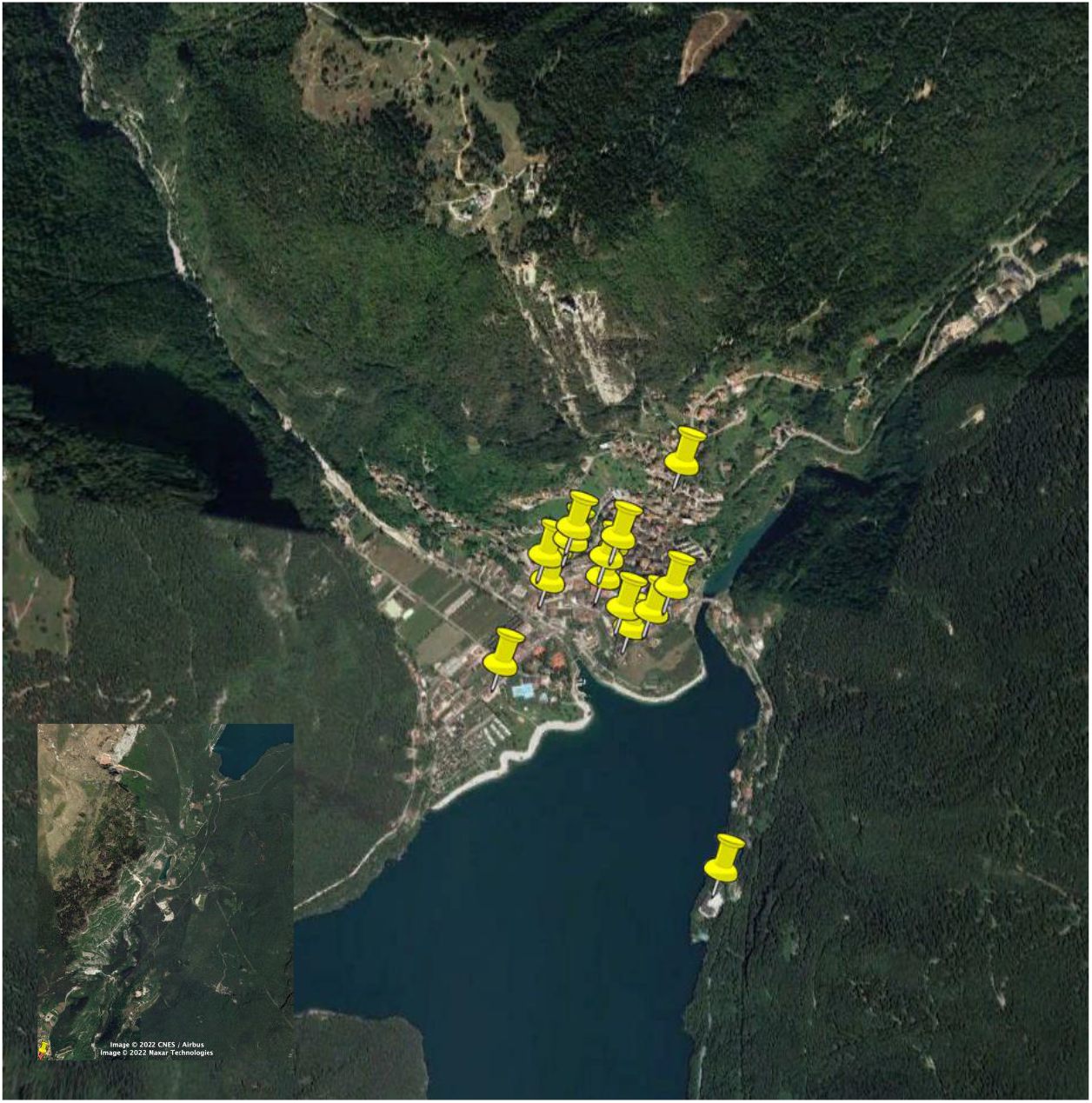

In September 2011, a previously healthy 70-year-old German tourist, fell ill with non-specific respiratory symptoms on the day he returned home after having visited this town and staying in one of its hotels. The patient required hospitalization and due to the suspicion of LD a test for *Legionella* urinary antigen (Biotest-ELISA, BioRad, Munich) was performed and found positive. *L. pneumophila* sg 1 was isolated from a tracheal secretion sample and typed as monoclonal subgroup Philadelphia (MAb 3/1 positive) and sequence type (ST) 901. Despite intensive treatment, the patient developed multi-organ failure and died 46 days after hospitalization. The German authorities informed the Italian health authorities that this infection might be related to the patient’s stay at “hotel F” in Italy (Table 1). Therefore, the Italian local health officers immediately inspected the hotel where the patient had stayed for seven days and water samples were collected from warm (WW) and cold water (CW) system. This investigation detected the *Legionella pneumophila* causative strain in concentration of 1,100 CFU/L (CW) and 12,700 CFU/L (WW) in a room and 42 CFU/L (CW) and 13,000 CFU/L (WW) in shared bathrooms. Colonies from the hotel were then typed and they resulted to belong to subgroup Knoxville ST901. The CFU values were over the threshold recommended by the Italian Guidelines for Prevention and Control of Legionnaires’ Disease [5], therefore disinfection of the water system and long-term measures were recommended to the hotel manager (unpublished data).

A query of European SBT database found only seven ST901 *L. pneumophila* sg 1, subgroup Philadelphia, out of more than 14,092 entries, including three clinical (among these the German patient above described) and four environmental isolates from Italy. Moreover, the first isolation dated back to 1987 and was from a patient, which was part of a cluster of two cases that occurred among a small group of British tourists staying in Molveno in a different hotel from the one where the German tourist had been. In-depth retrospective analysis revealed that after the first cluster in 1987, other TALD cases, both single cases and clusters, have occurred up to 2021 in that town, involving 22 hotels and 61 cases (among foreign and Italian tourists) with a ratio male to female 2,2:1 and an average patient age of 66.7 years.

Since over the years the *Legionella* collection has been enriched with isolates from these hotels, the aim of this study was to investigate the environmental and human isolates obtained in the 1987-2019 period at genome level, trying to detect the molecular characteristics that made these strains so virulent, and appearing as an endemic clone for that town.

## Material and Methods

### Bacterial strains

Overall, 40 *L. pneumophila* sg 1 (Lp1) strains (two human and 38 environmental isolates collected between 1987 and 2019), from the collection of the National reference laboratory for *Legionella*, Rome, Italy, were analyzed (Table 1). The two human isolates came from a German and a British tourist, who visited Molveno in 2011 and 1988, respectively. A third human sample, the DNA extracted from the EULV8733 Lp1 ST901 strain, isolated in 1987, was provided by the Public Health England. The environmental isolates were from 16 of the 22 hotels of Molveno, as shown in Figure 1.

### Culture, typing and DNA extraction of Lp1 isolates

Lp1 strains stored at −80°C were thawed and cultured on Buffered Charcoal Yeast Extract (BCYE, Oxoid) agar plates at 37°C for 48 h. A suspension in 1% formalin was used for MAb typing tests by indirect immune-fluorescence according to the Dresden panel [6]. Isolated colonies were suspended in PCR-grade water for the following automated DNA extraction using QIAcube and QIAmp mini kit (QIAGEN). The DNA extracts were used for SBT, performed as previously described [7-8], and for whole genome sequencing.

### WGS and assembly

Sequencing libraries were prepared using the NextEra XT library prep kit and then run on NextSeq-500 sequencer (Illumina, CA, USA): a 150-bp paired-end sequencing run was performed using a Mid output Kit v2. The fastq files were trimmed and assembled using the fully automated pipeline INNUca by Galaxy ARIES platform (https://aries-iss-it.iss.idm.oclc.org/) [9]. Genomes were annotated using Prokka [10]. All genome sequences were submitted to GenBank (Bioproject PRJNA931258; accession number are listed in Table1).

### cgMLST analysis

chewBBACA software [11] was used to convert the cgMLST scheme defined by Moran-Gilad et al (2015), which included 1,521 core genes [12]. The allelic profiles were then corrected to include only the subset of genes that were present in all the strains. The allelic profile output was then used to create minimum spanning trees with PHYLOViZ Online software [13].

### SNP analysis

Snippy software v4.5.0 was used to identify core SNPs (the SNPs located in core sites, genomic positions present in all the samples) on WGS data of each isolate. For each sequenced sample, the SNP calling was done aligning the reads to the contig assembly of sample 1862A (taken as reference). The SNP calling result folders were submitted to a second snippy run, giving the command “snippy -core”, that selected the core SNPs. The obtained core SNP alignment was used to construct phylogenetic trees by Phyloviz online software. To investigate if evidences of recombination events can be detected from SNPs, sequence reads were mapped to the CP013742 *Legionella* reference genome using Snippy v4.5.0, and horizontally transmitted SNPs were removed by Gubbins analysis [https://aries.iss.it/; 9].

Recombination in ST901 dataset was also investigated by alignment of two representative ST901 genomes versus other reference genomes (Table S1) using MAUVE program. The core genome regions, extracted with the command “strip_subset_LCBs” from the obtained alignment, were then analyzed for recombination events with RDP4 software using algorithm GENECONV (default parameters) and manual distance plot (step=200bps - window=600bps).

### Pangenome

Pangenome analyses were performed on genomes annotated by Prokka [9] by using Roary [15] with default values of parameters. The phylogenetic tree used as input for Roary was based on core SNPs and was calculated using Parsnp [17]. In order to characterize genes of the accessory genome islands, the amino-acidic gene sequences were submitted to a BLASTp search on the complete NCBI database (https://blast.ncbi.nlm.nih.gov/Blast.cgi), as well as to a resistome search on the Comprehensive Antibiotic Resistance Database (CARD - https://card.mcmaster.ca/) using the online tool Resistance Gene Identifier (RGI - https://card.mcmaster.ca/analyze/rgi). to identify genes possibly involved in resistance mechanisms. Blast atlas analysis by https://server.gview.ca/ was used to compare ST901 data set genomes.

## Results

### Typing and whole genome sequencing

The data set samples were typed as MAb 3/1 positive ST901 (allelic profile: 2,1,18,15,2,10,6) with the exception of the genome with ID 2016 that is a two loci variant of ST901 (ST1239), in alleles *fla*A and *mip* (i.e. 26 instead of 2 and 60 instead of 15, respectively) and it was isolated in only the “hotel N”.

The whole genome sequence of 41 Lp1 genomes, 38 environmental and three human, isolated following recurrent LD cases (n=61, all travel-associated) occurred in Molveno of the Northern Italy from 1987 to 2019 in 16 hotels, was determined. The genome sequences are approximately 3.5Kbp and the average G + C content is 38.46% (Table 1).

Prokka analysis revealed that all the ST901/1239 contain type I-C clustered regularly interspaced short palindromic repeats (CRISPR) and seven CRISPR-associated (cas) genes system (*cas*1/2/3/4/5c/7c/8c), as in *L. pneumophila* str. Toronto-2005, and a downstream CRISPR array including 32 spacers interlaced with the repeat sequence 5’-gtcgcgccccgtgcgggcgcgtggattgaaac-3’. Each *cas* gene showed 100% similarity with the corresponding genes in str. Toronto-2005.

Additionally, a 59,909 bp long contig was observed in all the genomes of the dataset. This appears a putative plasmid, as hinted by a sequence of approximately 80nt repeated at either ends, that is 98% similar to the *Legionella pneumophila* str. Lens pLPL plasmid (accession number: NC_006366). The contig contains 54 open reading frames with the exception of that found in the genomes 736A, 739A and 1152A, which has 53 open reading frames. Prokka analysis revealed that the 59,909 bp-contig also contains type I-F CRISPR-cas system characterized by five (cas1/cas3/csy2/csy3/cas6f-csy4 in the genomes with ID: 734A, 743A, 1026 and 2017) or six CRISPR-associated proteins (cas1/cas3/csy1/csy2/csy3/cas6f-csy4 found in the remaining genomes) and a downstream CRISPR array including 54 spacers interspaced with the palindromic repeat 5’- gttcactgccgtacaggcagcttagaaa-3’. In the 744A genome the CRISPR array is similar but consists of 50 spacers and additionally has four palindromic repeats with 3 more nucleotides (5’- tttctaagctgcctgtacggcagtgaac<u>tta</u>-3’, here underlined). Blast analysis revealed that the CRISPR-cas associated proteins have 100% identity to the I-F CRISPR-cas system found in *L. pneumophila* str Lens and spacers are the same in all the genomes of the ST901 data set. Among the 54 open reading frames annotated by Prokka, mainly represented by hypothetical proteins, there were *tra*C, *tra*N and *tra*D, *csr*A, *lex*A, *umu*C and *rec*D genes.

### cgMLST

The 41-genome ST901/ST1239 collection was de novo assembled and phylogenetic relationships inferred. cgMLST profiles, based on the subset of 1,453 cgMLST targets of the previously published 1,521 core gene scheme [10], were determined and compared with cgMLST profiles of 19 other genomes belong to other STs (Table S1), as shown in Figure 2. The minimum spanning tree groups the 41-genome collection in a unique clade with very few difference loci (one to seven) while the nearest ST23 shows 967 difference loci, highlighting a strict relationship among the ST901 and a considerable distance of ST901 from other STs (Figure 2).

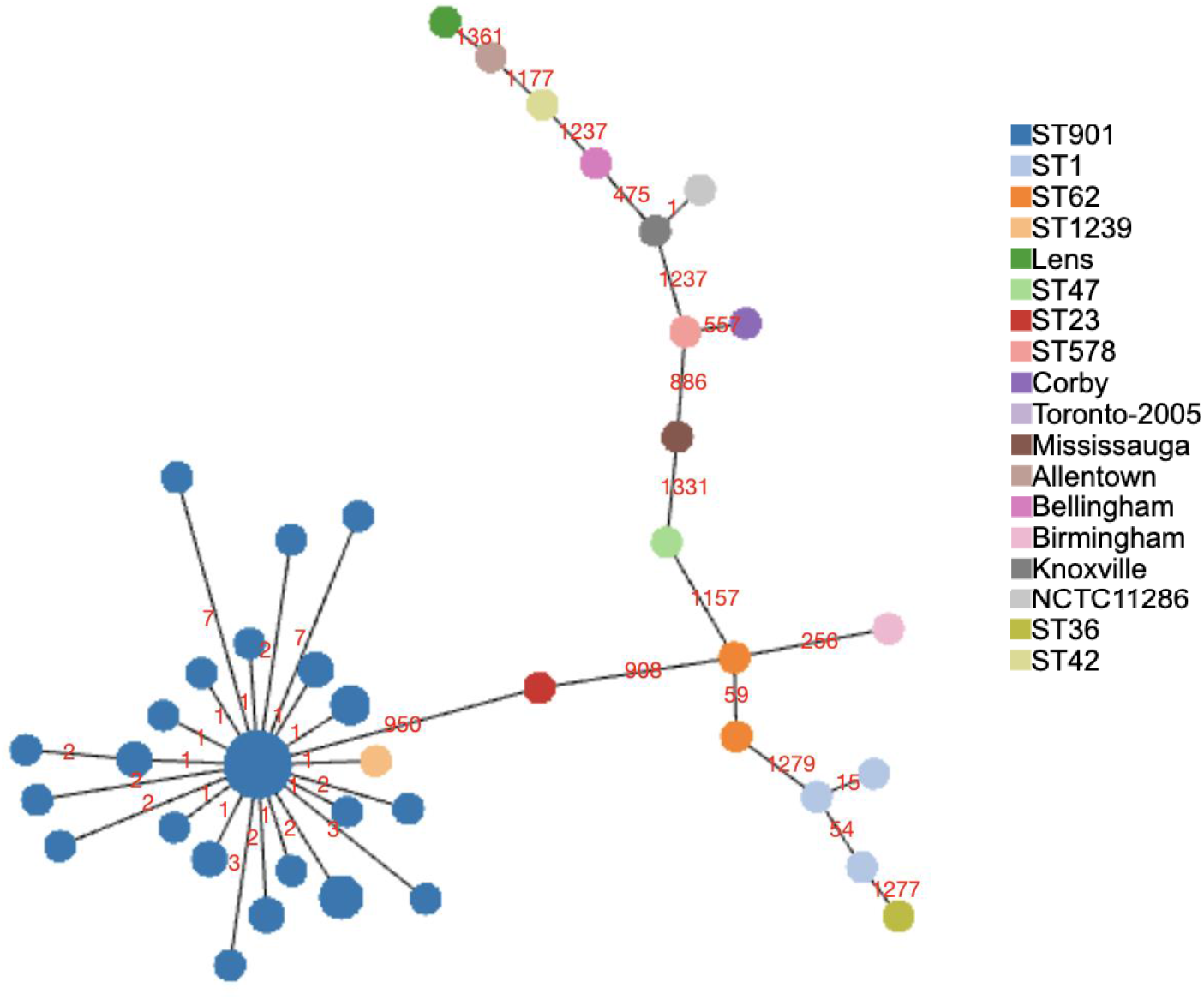

### SNPs analysis

SNPs analysis on the 41 ST901/ST1239 was also performed and Figure 3 represents the minimum spanning tree of the 68 core SNPs found in the ST901 data set, with maximum pairwise SNP differences ranging from 1 to 16, and with the average pairwise SNP differences of 5.18 (Table S2). The same analysis was repeated including the genomes of *L. pneumophila* str. Corby, Lens, Lorraine, Paris and ST23 and a maximum likelihood dendrogram based on parsimony tree highlighted the distance of the ST901 data set from these other genomes (Figure 3B, Table S1).

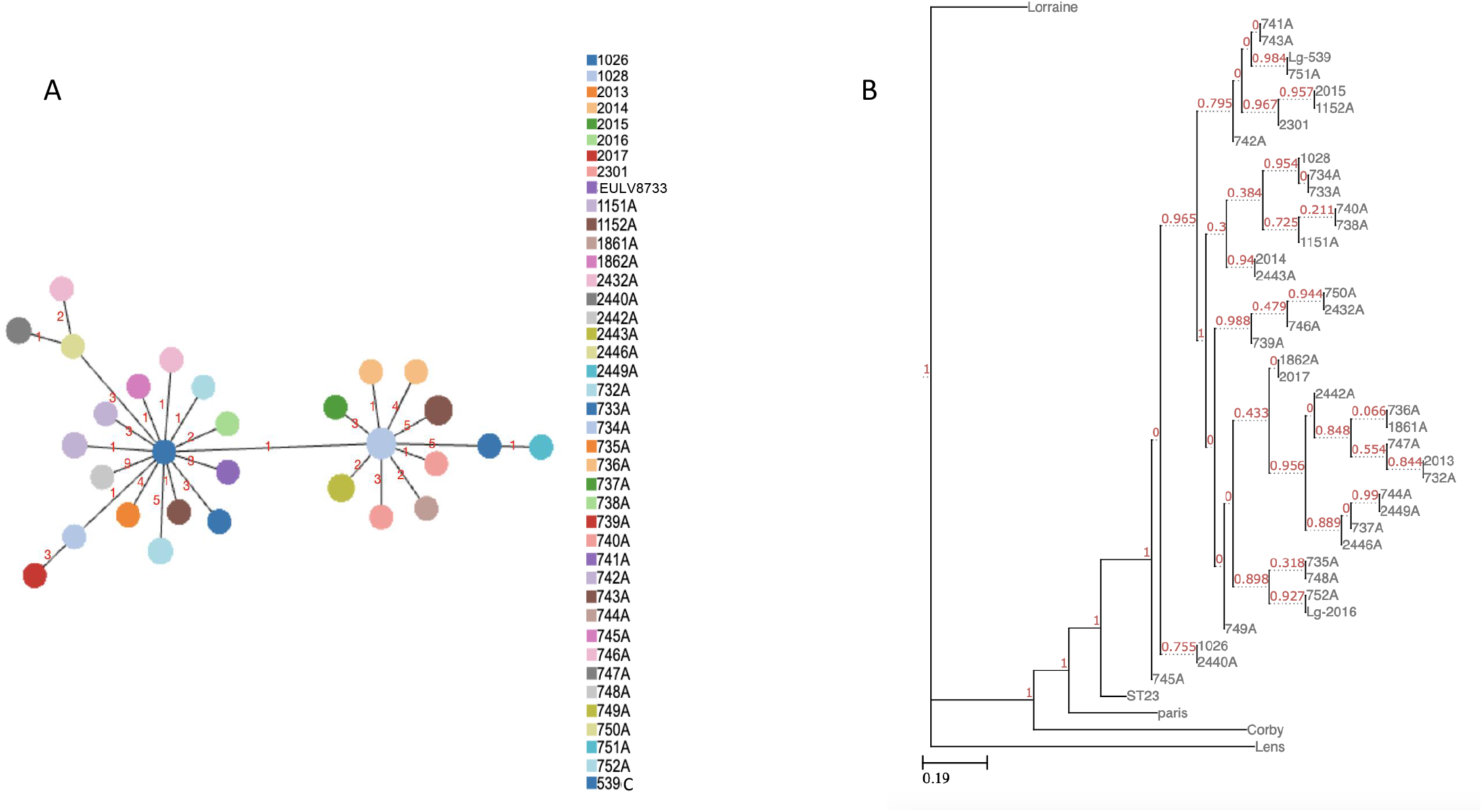

To investigate the phylogenetic relations on basis of vertical heredity and to check signals of recombination of ST901 strains among them and with other *L. pneumophila* reference genomes, Gubbins analysis was performed. Firstly, ST901 strains were compared to a reference genome (Genbank acc. n. CP013742) revealing that SNPs due to recombination range from 0 to 47.8% (a mean of 8,53% of the 3Mb genomes), while removing the recombined regions, to leave only the SNPs vertically transmitted, the maximum number of pairwise SNP differences was 33 with a mean of 11.28 of all ST901 (Figure S1).

Secondly, due to the high similarity of ST901 dataset, only the four ST901 subset, including 732A, 736A, 751A and 1862A, and a greater number of reference genomes (Table S1) were compared. The final cladogram-tree showed that ST901 are phylogenetically closer to ST23, Toronto_2005, Mississauga, Birmingham, ST62 and Pontiac genomes (Figure 4).

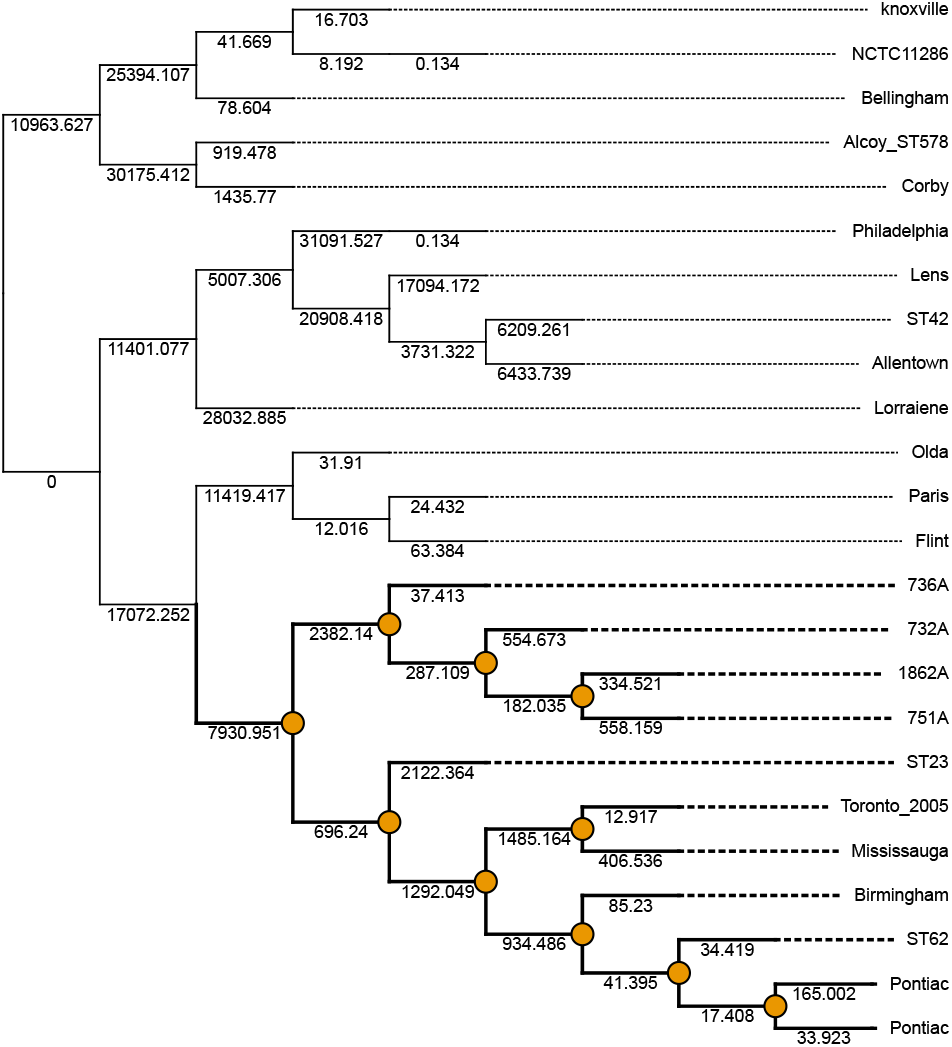

RDP4 software allowed to identify a region of approximately 200kb of ST901 genomes showing 100% identity to Lorraine sequence (from nt 273,749 to 530,163 on FQ958210 genome), in contrast with the rest of the Lorraine genome that poses this strain phylogenetically far from the ST901 dataset. Such region contains 212 genes among which there are ribosome genes, genes encoding for substrates of Dot/Icm secretion system, genes encoding for the components icm/dot of the secretion system (Table S3, Figure S2).

### Pangenome

The pangenome of the ST901/1239 data set consists of 3,116 core and 298 accessory genes out of a total of 3,476 genes, including 24 soft-core genes and 38 shell genes not counted in the core and accessory genomes. Taking as reference one of the genomes of the ST901/1239 dataset, blast atlas analysis revealed that the core genes of the ST901/1239 show an identity ranging from 80 to 100%. As shown in Figure 5A, accessory genes were mostly found (227 out of 298) in only the two environmental strains (ID 751A n=175 and 1862A n=52) that showed additional genomic regions. These additional regions 48,353bp (in 1862A) and 99,018bp (in 751A) in length, respectively, missing in all the other ST901 showed 100% identity with *L. anisa* isolate UMCG-3A plasmid p3A2 (accession number: NZ_CP029565) and 98% identity with *Legionella* sp. PC1000 plasmid pPC1000_1 (accession number: NZ_CP059403), respectively.

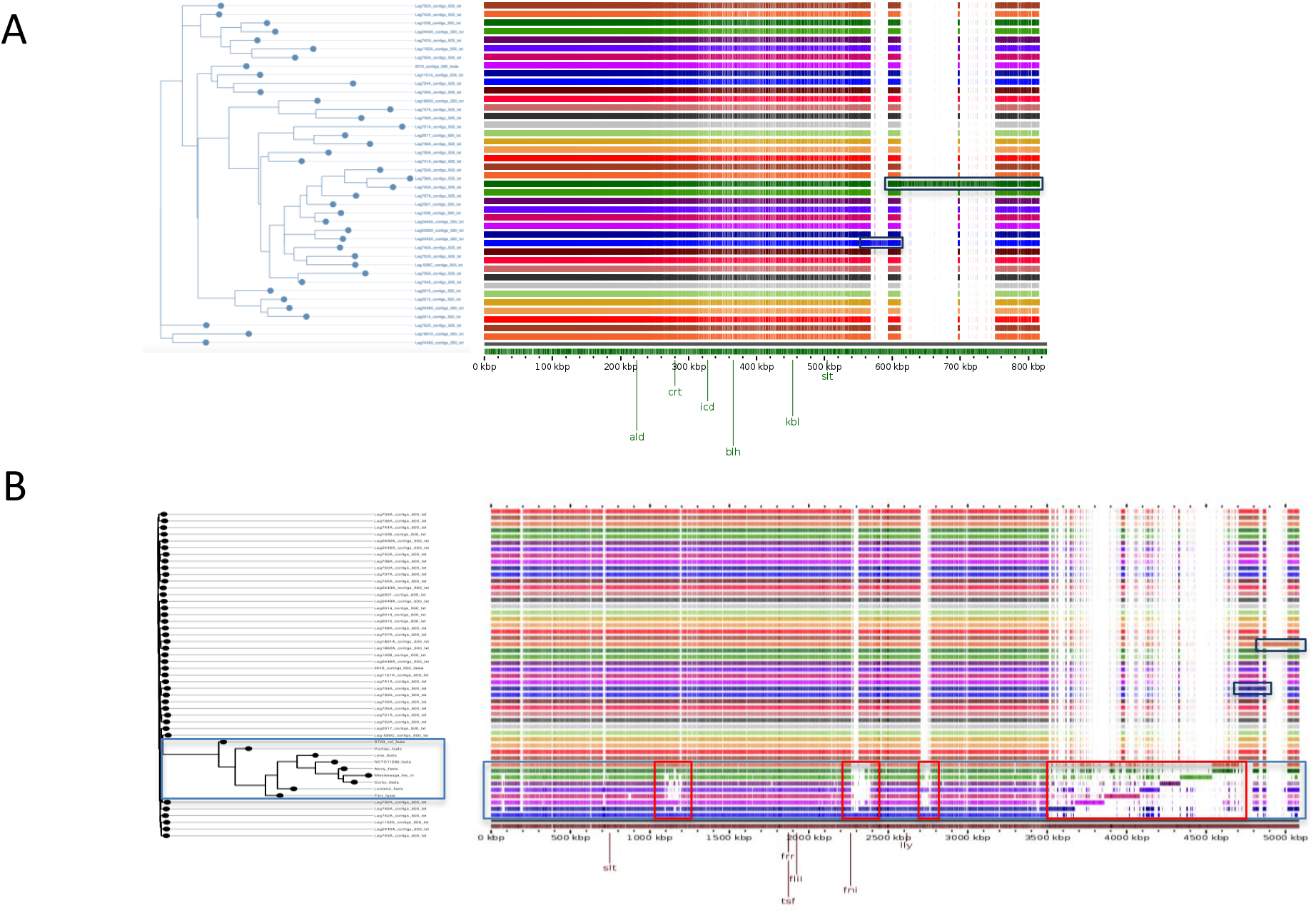

Additionally, in order to investigate similarities and differences in gene content with other unrelated genomes, the pangenome analysis was performed also including the following *L. pneumophila* genomes: Alcoy, Corby, Flint, Lens, Lorraine, Mississauga, Pontiac, NCTC 11286, EUL028-ST23. The obtained pangenome consisted of 6,050 genes, including 2,186 core genes and 3,864 accessory genes (counting also the soft-core genes n= 254 and the shell genes n= 745) and it is shown in Figure 5B. 36% of the pangenome was shared by all the genomes analysed (number of genes =2,186) while only 2% (number of genes = 122) of the pangenome was represented by genes specifically found in the ST901. Compared to the other genomes, the ST901 dataset and the EUL028 genomes (ST23) shared the highest number of accessory genes (n=676), while the lowest one is found between ST901 and str. Lens (n=270). As shown in Table 2, pangenome analysis allowed to verify that most of the accessory genes lie in accessory genome islands (AGIs) as already described by D’Auria et al [17]. Of the AGIs characterized by D’Auria et al., those resulting totally present in ST901 are TS1 (a transport and secretion island - also found in the strains Alcoy and Corby), C1 (the already quoted CRISP system I-F) and ND3 (an AGI with no clear role - also present in the strain Paris), while partially present are R1 (a resistance-related AGI - 13 genes out of 22 - present in most of the strains analyzed in the study), DT3 (DNA transfer-related AGI - 14 out of 45 - also found in Philadelphuia, Lens and Paris), PR1 (a phage related AGI - 9 out of 39 - also found in Philadelphia, Alcoy and Corby) and ND10 (another AGI with no clear role 22 out of 61- also present in the strain Paris). Additionally, pangenome analysis allowed to discover other AGIs not previously described, present specifically in the ST901 dataset or in ST901 and few other genomes, (Table S2). In particular, AGI_1, consisting of six genes encoding transposases/integrases, oxidoreductase, helix-turn-helix, and AGI_2, consisting of 16 genes including transposases, a family of proteins annotated as ATP-dependent Lon proteases, DNA repair proteins and several uncharacterized proteins, were found in Mississauga strain; AGI_3, with 30 genes, includes also transposases, conjugal transfer proteins, DNA repair proteins, thermonuclease family protein, metabolic enzymes, and some of these genes were also found in D-4959, Lens, Mississauga and ST62 strains; AGI_4 with six genes also found in few some other genomes; AGI_5 contains 17 genes encoding integrases, YfjI family protein, S-adenosylmethionine protein families, helix-turn-helix transcriptional regulator, and it appears specifically found in ST901 dataset; AGI_6 with six genes also found in few other genomes; AGI_7 contains numerous genes with homology to resistance genes, and overall it includes 46 genes among which integrases, metabolic enzymes, LuxR transcriptional regulator, heat-shock protein, aminoglycoside phosphotransferase family proteins, many GNAT family N-acetyltransferases, type IV secretion system effectors and it was in part found also in Birmingham, Pontiac, ST23 and ST62 strains; AGI_8 with 21 genes encoding metabolic enzymes, integrases, AAA family ATPase, LysR family transcriptional regulator, and in part found in Birmingham, Pontiac, ST23 and ST62 strains; AGI_9, with only three genes one of which also found in NCTC12273 strain; AGI_10 with the seven genes corresponding to the type I C-CRISPR-cas system, described above; AGI_11 exclusively found in ST901 dataset; AGI_12 with metabolic enzymes and integrase was found also in NCTC11985 and Pontiac genomes; AGI_13 and AGI_14 of three and two genes; AGI_15 and AGI_16 with 12 and three genes respectively, were found only in ST901 dataset. As already observed by D’Auria et al [17], the accessory genes of the ST901 dataset were prevalently hypothetical proteins (71%).

## Discussion

To our knowledge ST901 have been isolated isolated only in Molveno, a little town of the Northern Italy, causing recurrent LD cases over a 32-year period and until now it appears to be localized exclusively in that town. Due to the large number of TALD sporadic cases and clusters among possibly healthy people, without a space-temporal context of an outbreak, ST901 were analysed and compared with genomes of other unrelated Legionella strains with regard for specific virulence traits.

Noteworthy is the presence in the ST901/1239 data set of type I-C and I-F CRISPR-cas systems, the first localized on the chromosome and the other one on a putative plasmid, respectively. CRISPR-cas systems are multiple adaptive immune systems, present in most archea and bacteria, that during the adaptation stage incorporate fragments of foreign DNA, providing sequence-specific protection from invading viruses and plasmids [18]. These fragments of foreign DNA are known as protospacers and they constitute the CRISPR array as new spacers. The type I-C CRISPR-cas system found in ST901 has been previously described in the ST222 Toronto-2005 strains, which caused an explosive LD outbreak in Toronto, and in other four ST222: Toronto-2000, Ottawa-2005, Mississauga-2006 and London-2007 [19-20]. As discussed later, a possible phylogenetic relation between ST901 and ST222 was highlighted also by pangenome analysis. The I-C CRISPR-cas system has been described to be active and adaptive, with a CRISPR array constantly evolving by acquiring new spacers upstream of ancestral. The CRISPR array of ST901 dataset consists of primed spacers, suggesting that this data set has been invaded by the same invaders, and the observed mismatches in both protospacers and their flanking regions indicate an actual stimulation to spacer acquisition, making bacteria able to resist to phage or virus that escape the interference of older spacers (data not shown; 21). Additionally, the ST901 data set is characterized by the type I-F CRISPR-cas system with 100% identity to the type I-F systems that are present on annotated plasmids, strain Lens pLPL and strain Mississauga-2006. This let suppose that most probably the I-F system of the ST901 dataset was horizontally acquired, being located on a putative plasmid [22]. The co-occurrence with the chromosomal type I-C system has been already demonstrated for strains Mississauga-2006 and FJAD01, letting suppose that this is not a so rare event in *L. pneumophila* isolates [23]. Considering the long stay in the environments where the ST901 were isolated, the co-occurrence of type I-C and I-F systems most likely contributed to maintaining a potent adaptive immune response to invading DNA, ensuring survival in the environment.

cgMLST, SNP and pangenome analyses demonstrated that this genome collection, including the genome with a slightly different allelic profile that determines the ST1239, independently on the year of isolation, are highly similar but not closely related to other STs. In contrast to the phylogenetic relations established on basis of the whole genome, an approximately 200Kb long region, encoding about 200 genes among which the components icm/dot of the type IVB secretion system, shows an high similarity (nearly 100%) with Lorraine strain, also known as ST47, probably as result of a homolog recombination event. The Lorraine ST47, firstly described in 2008, nearly exclusively isolated in North-west Europe, is characterized by a highly conserved core genome and a diverse accessory genome and no specific virulence related genes which explained its high disease potential [24-26]. Although no recombination was detected within the ST47 lineage [26], it could be possible that ancestrally it have recombined with other Legionella strains and also with ST901. Additionally, pangenome analysis demonstrated that when genomes other than ST901 were included, only the 2% of the genes were typically found in ST901/1239 dataset and the ST23, Toronto_2005, Mississauga, Birmingham, ST62 and Pontiac genomes, albeit at some distance, were found to be phylogenetically closer to ST901 data set. Indeed, this analysis allowed also to observe that most of the accessory genes are organized in islands and many of these islands shared the same genes as the islands previously observed by D’Auria et al. in the Alcoy, Paris, Lens, Corby and Philadelphia genomes, such as resistance-related islands, transport/secretion systems, DNA transfer-related and phage-related islands [17]. The occurrence of genomic islands related to virulence as those found in genomes believed so virulent as Alcoy and Corby strains, as well as the presence of CRISPR-cas islands could explain the major virulence for humans and a great capacity of persistence in the environment of the ST901 dataset compared to other *Legionella* strains. About the AGIs found in the ST901 dataset, almost all the genes showed 100% identity with the same genes in other genomes. AGI_7 and AGI_8, characterized by several genes encoding type IV secretion system effectors, showed 100% identity with the genes of ST23, Birmingham, ST62 and Pontiac where these AGI were also observed. In particular, AGI_7 is characterized by numerous genes showing homology to resistance genes that would let suppose being a potential resistance-related island.

A lot of the AGIs contain transposases/integrases however, among those specifically found in ST901, only few genes with homology to virulence factors were detected. Four of the 16 accessory genome islands (AGI_5, 11, 14 and 16) appear to be strain-specific. In particular AGI_5 contains the proteins HxsC, B and D characterized by the repeats His-Xaa-Ser that appear to be part of a peptide modification system. In general, proteins flanking His-Xaa-Ser systems lack pairwise homology to each other, suggesting that the His-Xaa-Ser system does indicate the presence of any one specific type of mobile element [27].

Unfortunately, data on *Legionella* strains isolated from private homes or other kind of buildings in Molveno were not available. However, epidemiological surveillance data indicate the absence of LD cases among the residents in Molveno, this suggests that the particular virulence of ST901 and factors related to the characteristics of the hotel water systems are responsible for the high number TALD cases in this town. As reported by Stout et al [28] in contrast to private homes, hotel water systems are often characterized by the presence of tanks with temperature stratification, no operation periods, recirculating system seeding the bacteria to distal site, etc., favouring *Legionella* proliferation when no appropriate maintenance is carried out. Once a hotel water system is contaminated by *Legionella*, the likelihood to have LD cases is expected to increase as the size of the hotels, as well as the age of the clients and the length of stay [29].

Besides, investigations carried out on the possible contamination of the two water sources that feed the municipal water of Molveno, gave always negative results (unpublished data).

In conclusion, the genome analysis of the ST901 dataset demonstrated that these genomes have maintained their setup over time, which most probably allowed them causing numerous TALD cases. ST901 appear to share specific virulence related traits with some of the most virulent *Legionella* strains that could be possible descent. Their success in the environment could be due to the capacity to persist in the water system, thanks to the co-occurrence of two phage resistance systems. The presence of virulence related AGIs could have made these strains enough virulent to cause such a high number of TALD cases.

## Acknowledgements

The authors are grateful to Dr. Italo Dell’Eva and Dr. Fabrizia Helfer of the Trento Public Health Laboratory for providing them with the environmental strains analyzed in this study collected from the water system of Molveno accommodation facilities, as well as Dr. Franco Guizzardi of the Provincial Agency for Health Services of the Autonomous Province of Trento for his valuable collaboration in the epidemiological and environmental investigations carried out following the LD cases.

The authors wish to thank the Saxony health service in Dresden for support. We thank Kerstin Lück, Susann Menzel (Dresden) Teresa Stocki and Massimo Mentasti (London) for their excellent technical assistance.

